# A novel bacterial protease inhibitor adjuvant in RBD-based COVID-19 vaccine formulations increases neutralizing antibodies, specific germinal center B cells and confers protection against SARS-CoV-2 infection

**DOI:** 10.1101/2021.12.07.471590

**Authors:** Lorena M. Coria, Lucas M. Saposnik, Celeste Pueblas Castro, Eliana F. Castro, Laura A. Bruno, William B. Stone, Paula S. Pérez, M. Laura Darriba, Lucia B. Chemes, Julieta Alcain, Ignacio Mazzitelli, Augusto Varese, Melina Salvatori, Albert J. Auguste, Diego E Álvarez, Karina A. Pasquevich, Juliana Cassataro

## Abstract

In this work we evaluated recombinant receptor binding domain (RBD) based vaccine formulation prototypes with potential for further clinical development. We assessed different formulations containing RBD plus Alum, AddaS03, AddaVax or the combination of Alum and U-Omp19: a novel *Brucella* spp. protease inhibitor vaccine adjuvant. Results show that the vaccine formulation composed of U-Omp19 and Alum as adjuvants have a better performance: it significantly increased mucosal and systemic neutralizing antibodies in comparison to antigen plus Alum, AddaVax or AddaS03. Antibodies induced with the formulation containing U-Omp19 not only increased their neutralization capacity against the wild-type virus but also cross neutralized alpha, lambda and gamma variants with similar potency. Also, addition of U-Omp19 to vaccine formulation increased the frequency of RBD-specific geminal center B cells and plasmablasts. Additionally, U-Omp19+Alum formulation induced RBD-specific Th1 and CD8^+^ T cell responses in spleens and lungs. Finally, this vaccine formulation conferred protection against an intranasal SARS-CoV-2 challenge of K18-hACE2 mice.

## Introduction

Severe acute respiratory syndrome coronavirus 2 (SARS-CoV-2) is the causative agent of coronavirus disease 2019 (COVID-19) that developed into a global pandemic causing (as of November 30, 2021) over 260 million cases and over 5.2 million deaths worldwide (Weekly epidemiological update, World Health Organization, WHO). Mass vaccination offers the most efficient public health intervention to control the pandemic. Several vaccines have been shown to be effective and have been either approved or authorized for emergency use in different countries (Status of COVID-19 Vaccines, WHO).

Despite the efforts made to vaccinate people, it is still too early to establish the durability and extent of protection, and recent data on approved vaccines have showed a diminished efficacy six months after vaccination^1-3^. Most importantly, it is critical to find a way to optimize the existing vaccines to protect against the prevalent SARS-CoV-2 variants of concern (VOC) that are spreading globally ^4^. Evidence of waning immunity and viral diversification create a possible need for a booster vaccine dose to protect the population^5^ leading advisory health agencies to recommend and additional dose of a COVID-19 vaccine. For all these reasons, there is a need to produce safer, more effective, highly scalable, and more affordable COVID-19 vaccines locally or regionally.

Most of the approved vaccines are mRNA-based, vector-based or inactivated viruses. Currently, there are a few protein-based subunit vaccine candidates in late phase trials^6^. Subunit vaccines are a well-known platform, and many subunit vaccines are already in widespread use. Protein subunit vaccines are easy to produce and safe, but in practice, they require a suitable adjuvant to stimulate the host immune response.

Subunit vaccine candidates in development are mainly based on Spike protein or the SARS-CoV-2 receptor-binding domain (RBD). RBD is located within the S1 subunit of the Spike. Angiotensin converting enzyme 2 (ACE2) is the functional receptor for SARS-CoV-2 comprising a critical factor for SARS-CoV-2 to enter into target cells and RBD is a key functional component that is responsible for binding of SARS-CoV-2 to host cells^7,8^. It is therefore not surprising that antibodies directed against the RBD or overlapping with the ACE2 binding region are strongly neutralizing, making the RBD a promising subunit vaccine candidate^9^. RBD-based antigens have been described in previous studies for SARS-CoV and MERS-CoV vaccine development^9,10^. RBD from SARS-CoV-2 is an ideal antigen for vaccine formulations because of its high expression levels, ease of manufacturing, stability, and capacity to elicit functional antibodies ^11^.

Although there is not a defined immune correlate of protection from SARS-CoV-2 infection yet, it has been proposed that neutralizing antibody levels are highly predictive of immune protection^12,13^. It has been found a strong correlation between vaccine-induced neutralizing antibodies (nAbs) and a reduction of viral loads in non-human primates and humans after SARS-CoV-2 infection^14,15^. T cell responses also play important protective roles in SARS-CoV-2 infection. The depletion of T cells in rhesus macaques has been shown to impair virus clearance^14^. In humans, virus-specific CD4^+^ and CD8^+^ T cell responses are associated with milder disease, indicating an involvement in protective immunity against COVID-19. Therefore, an ideal vaccine is expected to induce both the humoral and cellular arms of the immune system.

Vaccine adjuvants can enhance the magnitude, breadth, and durability of the immune response. Following its introduction in the 1920s, alum remained the only adjuvant licensed for human use for the next 70 years, however, five new adjuvants have been included in licensed vaccines until present^16^. The design and selection of adjuvants for COVID-19 vaccine formulations are key to induce optimal immune responses with adequate safety profiles. The introduction of novel adjuvants which have been shown to induce both humoral and cellular immune responses could be more favorable.

In previous works we demonstrated that a bacterial protease inhibitor from *Brucella abortus* (U-Omp19) can be used as an adjuvant in parenteral and oral vaccine formulations^17-20^. U-Omp19 parenteral delivery induces the recruitment of CD11c^+^ CD8α^+^ dendritic cells (DCs) and monocytes to lymph nodes where it partially limits *in vivo* antigen (Ag) proteolysis inside DCs and increases Ag intracellular half-life. Consequently, U-Omp19 enhances Ag cross-presentation by DCs to CD8^+^ T cells. Antitumor responses were elicited after U-Omp19 co-administration, increasing survival of mice in a murine melanoma challenge model. Moreover, subcutaneous, or intramuscular co-administration of U-Omp19 with *Trypanosoma cruzi* Ags conferred protection against virulent parasite challenge, reducing parasitemia and increasing mice survival^19,21^. When U-Omp19 was co-delivered orally it increased mucosal Th1, Th17, CD8 T and Ab responses and reduced parasite or bacterial loads after oral challenge with virulent *Toxoplasma gondii* or *Salmonella*^*20*^. Thus, U-Omp19 is a promising novel adjuvant able to promote specific Th1 and CD8^+^ T cell immune responses in addition to Ab responses.

Here, we present preclinical data of a COVID-19 recombinant RBD-based vaccine candidate formulated with Alum and U-Omp19 adjuvant with potential for further clinical development. Development of recombinant protein based COVID-19 vaccines could allow vaccine availability even in low-and middle-income countries at affordable costs.

## RESULTS

### Antigen expression and characterization

In this study, RBD was used as the vaccine antigen. A monomeric version of RBD preceded by SARS-CoV-2 spike signal peptide for secretion and a C-terminal hexahistidine (6xHis)-Tag was expressed after plasmid transfection in HEK-293 cells. The RBD segment selected in this study (residues 319-341) contains 8 predicted immunodominant CD4^+^ T cell epitopes and 17 predicted CD8^+^ T cell epitopes in addition to B cell epitope motifs^22^.

Recombinant RBD was purified from cell culture supernatant through a single-step Ni-NTA affinity chromatography. SARS-CoV-2 RBD protein had high expression with remarkable purity (**Fig. 1A**). Noteworthy, purified RBD was recognized by polyclonal antibodies in sera from a convalescent patient infected with SARS-CoV-2 (**Fig. 1B**). Analytical evaluation by size exclusion chromatography revealed that the recombinant protein is monodispersed. The sample eluted as a single peak with an apparent molecular weight of 40.3 KDa, representing >95% of the sample. (**Fig. 1C**). Endotoxin levels measured in the purified protein were ≤1,25EU per mg, this value is significantly lower than the maximum recommended endotoxin level for recombinant subunit human vaccines^23^.

**Figure 1.**
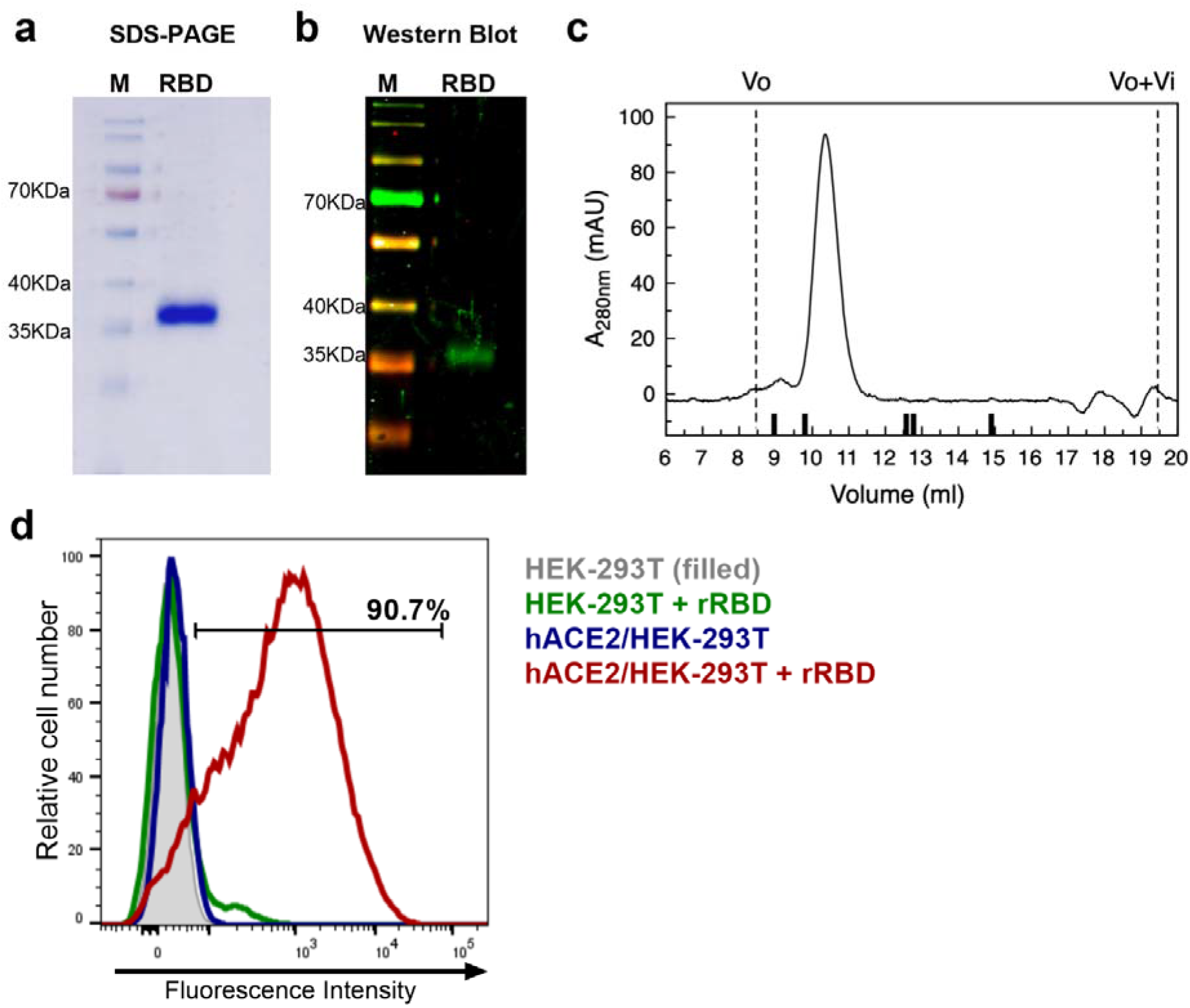
Characterization of recombinant RBD. **A**. Coomassie blue-SDS-PAGE stained of reduced RBD. M: page ruler. **B**. Western Blot analysis of recombinant RBD produced in HEK-293 cells using a convalescent human serum as primary antibody. M: page ruler. **C**. Representative size exclusion chromatography elution profile of recombinant RBD. Black bars represent MWM, from left to right: BSA (66.4 KDa), OVA (44.3 KDa), Papain (23 KDa), RNaseA (13.7 KDa), Aprotinin (6.5 KDa), Vo and Vo+Vi are shown as dashed lines. **D**. Binding of recombinant RBD (rRBD) to HEK-293T cells expressing hACE2. HEK-293T were used as control. Histograms and percentage of cells positive for RBD are shown.

Purified RBD was also assessed for its direct binding to the human ACE2 receptor in ACE2-expressing HEK-293T cells by flow cytometry. Strong binding of recombinant RBD to hACE2-HEK-293T cells was evidenced (>90%) (**Fig. 1D**). This result confirms that purified RBD binds to cell-associated hACE2 receptor suggesting that it has a correct folding.

### All vaccine formulations induce serum RBD specific IgG responses while only U-Omp19+RBD+Alum formulation induces specific IgA in BAL after intramuscular immunization

The immunogenicity of a variety of vaccine formulations comprising recombinant RBD with different adjuvants was evaluated in mice. We assessed formulations containing approved adjuvants for human use, including Aluminium hydroxide, AddaS03 (similar to MF59) or AddaVax (similar to AS03) and the combination of Alum and U-Omp19, a novel adjuvant developed in our laboratory that demonstrated vaccine adjuvant properties when co-administered with different Ags at pre-clinical stages.

Recombinant RBD was formulated with aluminium hydroxide (-Alum-Alhydrogel 2%) alone, Alum plus U-Omp19, AddaS03 or AddaVax. BALB/c mice received two doses at day 0 and 14 via intramuscular (i.m.) route (**Fig. 2A**). After the first dose, animals immunized with RBD+Alum or RBD+Alum+U-Omp19 produced a specific anti-RBD IgG response in serum (GMT at day 14: 4222 and 5572 respectively, **Fig. 2B**). Anti-RBD IgG titers increased after the second dose reaching a plateau (GMT at day 42: 215269 and 323050 respectively, **Fig. 2B**). The groups of animals immunized with formulations containing AddaVax or AddaS03 as adjuvants failed to induce specific antibodies after the first dose (**Fig. 2B**) but showed a significant anti-RBD IgG response after two doses (GMT at day 42: 337794 and 215269 respectively). RBD-specific IgG subclasses were evaluated one month after last immunization demonstrating that all vaccine formulations induced higher titers of IgG1 than IgG2a in serum of mice (**Fig. 2C**). Specific-IgA at the low respiratory tract has an important role to control virus dissemination^24^. Interestingly, levels of RBD-specific IgA in the bronchoalveolar lavage (BAL) of mice were higher in the group that received RBD+Alum+U-Omp19 than in groups that received AddaVax or AddaS03 (**Fig. 2D**). Besides, anti-RBD IgA was measured in serum samples revealing that groups containing Alum or Alum plus U-Omp19 as adjuvants induced significant higher levels compared to PBS or groups containing AddaVax or AddaS03 as adjuvants (**Fig. 2E**).

**Figure 2.**
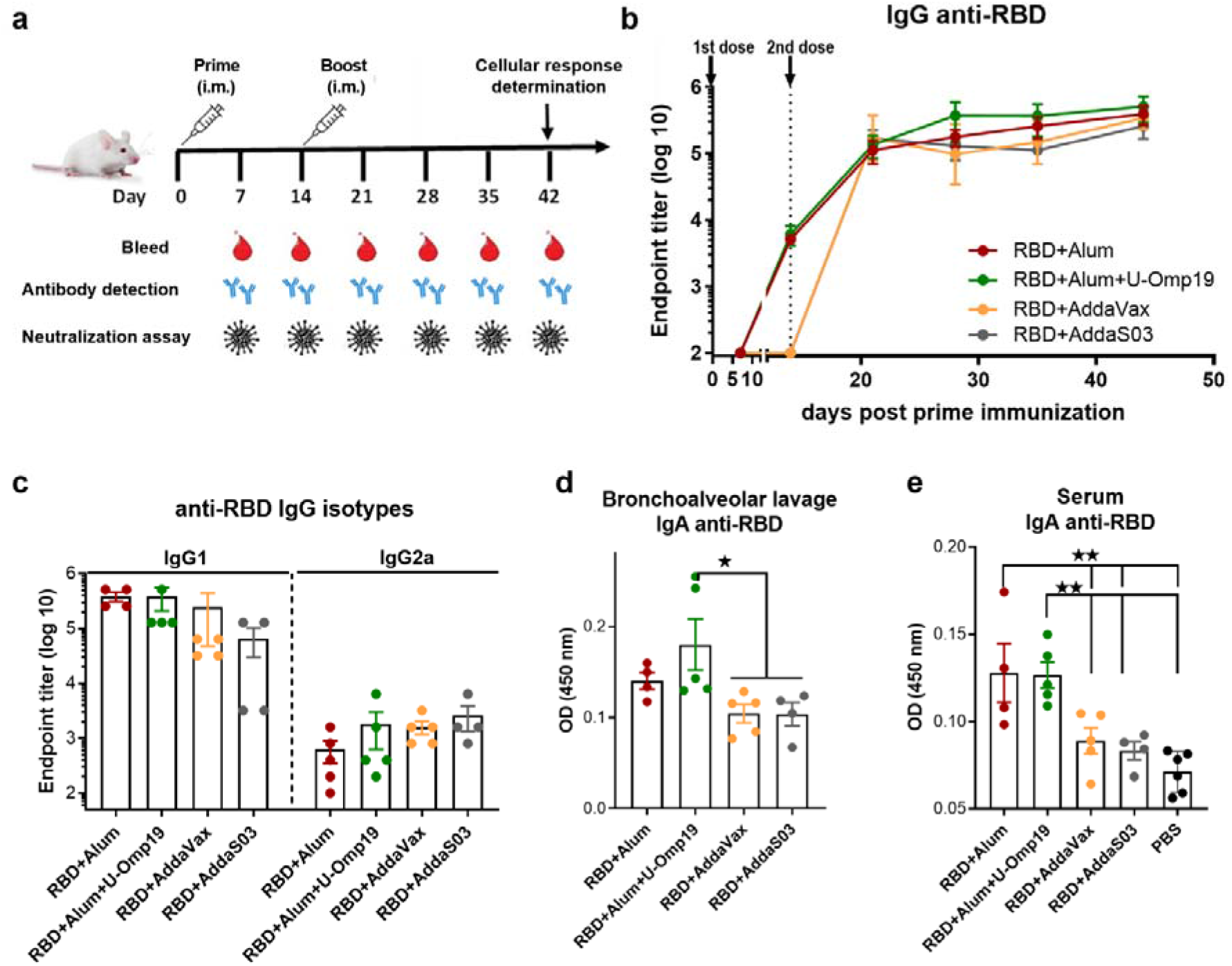
BALB/c mice immunized with RBD plus different adjuvants induced RBD-specific antibodies. **A**. Immunization protocol scheme. BALB/c mice were vaccinated at day 0 and day 14 via i.m. with: RBD+Alum (n=5), RBD+Alum+U-Omp19 (n=5), RBD+AddaVax (n=5) or RBD+AddaS03 (n=4). Serum samples were obtained at indicated time points for ELISA and neutralization assays. **B**. Kinetics of RBD-specific IgG endpoint titers in sera of immunized animals by ELISA. Points are means ± SEM. **C**. RBD-specific IgG subclasses (IgG1 and IgG2a) titers in sera of immunized animals at day 42 post prime immunization. Detection of RBD-specific IgG (**D**) and IgA (**E**) in the bronchoalveolar lavage of immunized mice at day 42 post prime immunization. Data are optical density (OD) at 450nm. *p<0.05. **p<0.01. Mann Whitney test.

### U-Omp19+Ag+Alum formulation significantly increases mucosal and systemic neutralizing Abs in comparison to Ag plus Alum, AddaVax or AddaS03

Next, the neutralization capacity of the vaccine induced antibodies was evaluated using an HIV-based pseudovirus neutralization assay (PsVNA). All vaccine formulations induced serum neutralizing antibodies against the SARS-CoV-2 spike pseudotyped virus (**Fig. 3A**). Remarkably, immunization with two doses of the formulation containing Alum plus U-Omp19 as adjuvants induced a ten-fold increase in the neutralization titer (GMT 325.1 95%CI 103.8-1018) compared to the groups immunized with Alum alone as adjuvant, AddaVax or AddaS03 (GMT 34.2 95%CI 3.790-308.5, **Fig. 3A**). This increment was statistically significant (*p*=0.0257 *vs* RBD+Alum, *p* =0.0259 *vs* AddaVax and *p* =0.022 *vs* AddaS03). AddaVax or AddaS03 as adjuvants induced a titer of neutralizing antibodies similar to Alum alone (**Fig. 3B**).

**Figure 3.**
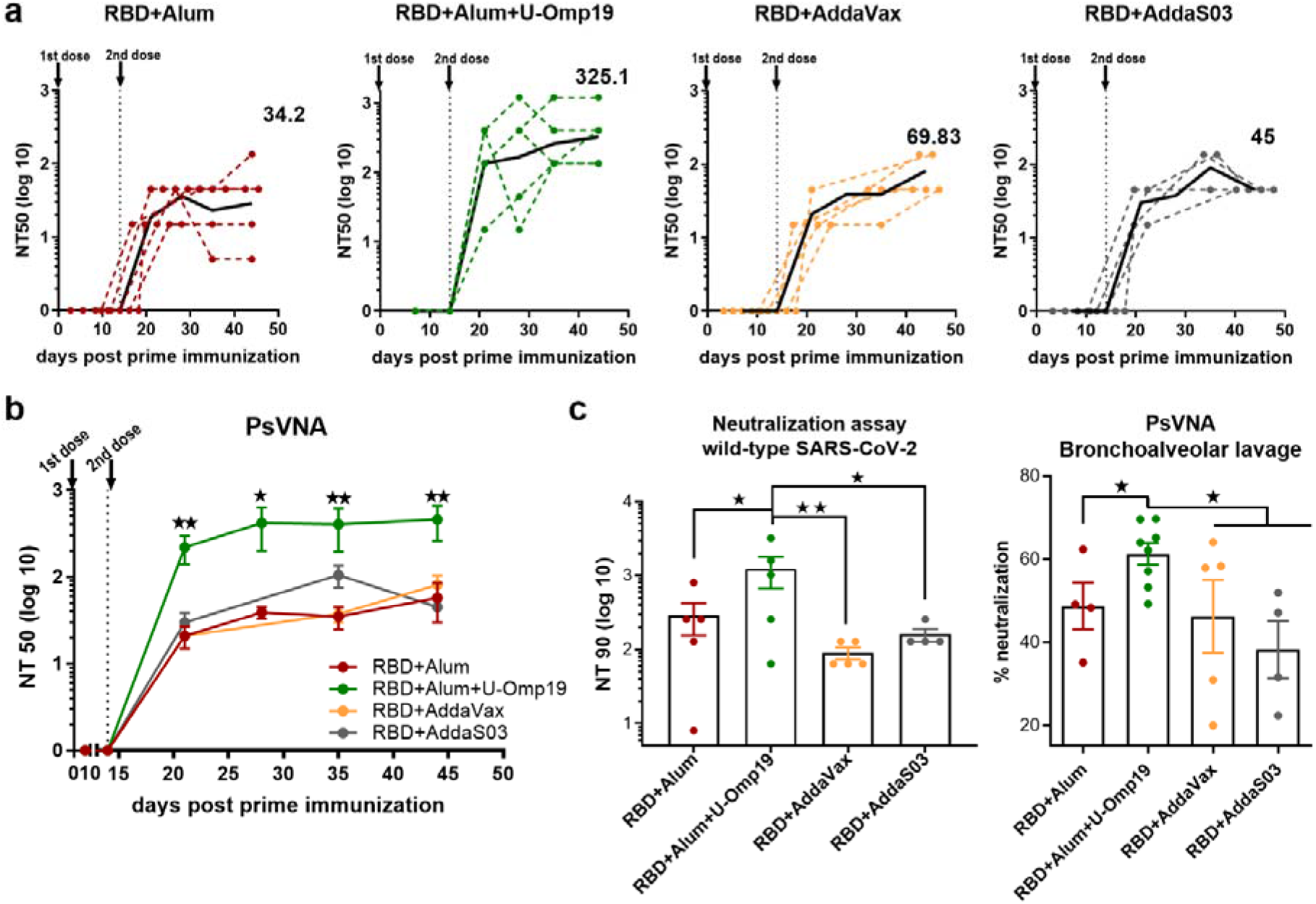
Immunization with RBD+Alum+U-Omp19 increases systemic and mucosal neutralizing antibody titers. **A**. BALB/c mice were vaccinated as described in Fig. 2. Serum neutralizing-antibody titers determined by pseudo-typed SARS-CoV-2 assay for each group of vaccinated mice at different time points. The black lines represent the geometric mean of all data points and numbers are GMT on day 42. Dotted lines represent the mean titer of each mouse at indicated time points. Titers correspond to the 50% of virus neutralization (NT50). **B**. Kinetics of neutralizing antibody titers of all groups determined by pseudo-typed SARS-CoV-2 assay. Points are Means ± SEM. Titers correspond to the 50% of virus neutralization (NT50). *p<0.05. **p<0.01. One way ANOVA with Bonferroni post-test. **C**. Neutralizing antibody titers against wild-type SARS-CoV-2 virus at day 42 post prime immunization. Neutralization titer was defined as the highest serum dilution without any cytopathic effect in replicable wells (NT 90). Data are shown as means ± SEM. *p<0.05. **p<0.01. One way ANOVA with Bonferroni post-test. **D**. Determination of neutralizing antibodies in the bronchoalveolar lavage by pseudo-typed SARS-CoV-2 assay at day 42. Data are expressed in percentage of neutralization compared with controls (virus alone). *p<0.05. T test.

To assess the functionality of vaccine-elicited antibodies against the wild-type SARS-CoV-2, neutralization assay with sera from immunized animals was performed. Similar to the results obtained using the pseudovirus system, one month after the second dose, RBD+Alum+U-Omp19 immunized mice had significant higher virus neutralization antibody titers in serum (**Fig. 3C**, GMT 612.1 95%CI 87.80-4267) than mice immunized with RBD+Alum (**Fig. 3C**, GMT 140 95%CI 16.34-1199) or plus commercial adjuvants (AddaVax or AddaS03). These results further confirm the data obtained by PsVNA.As SARS-CoV-2 initially infects the upper respiratory tract, its first interactions with the immune system must occur predominantly at the respiratory mucosal surfaces. Mucosal responses may be crucial to stop person to person transmission of this virus. Thus, examination of neutralizing activity in BAL was performed using pseudotyped virus system. Of note, adding U-Omp19 as an adjuvant to the formulation increased wild-type virus neutralization in the BAL of mice compared with the vaccine adjuvanted with Alum alone, AddaVax or AddaS03 (**Fig. 3D**, \**p*<0.05).

### Neutralizing antibodies last over 5-6 months after immunization with U-Omp19+Ag+Alum

Duration of vaccine immunity is key to estimate how long protection lasts. To this effect, we evaluated the level of total antibodies over 175 days after prime immunization with the vaccine formulation containing U-Omp19 and alum as adjuvants. Interestingly, titers of anti-RBD IgG antibodies remained stable at least 5-6 months after i.m. immunization of mice with this formulation (**Fig. 4A**). Remarkably, neutralizing capacity of the antibodies remained stable till day 175 post prime immunization (**Fig. 4B**)

**Figure 4.**
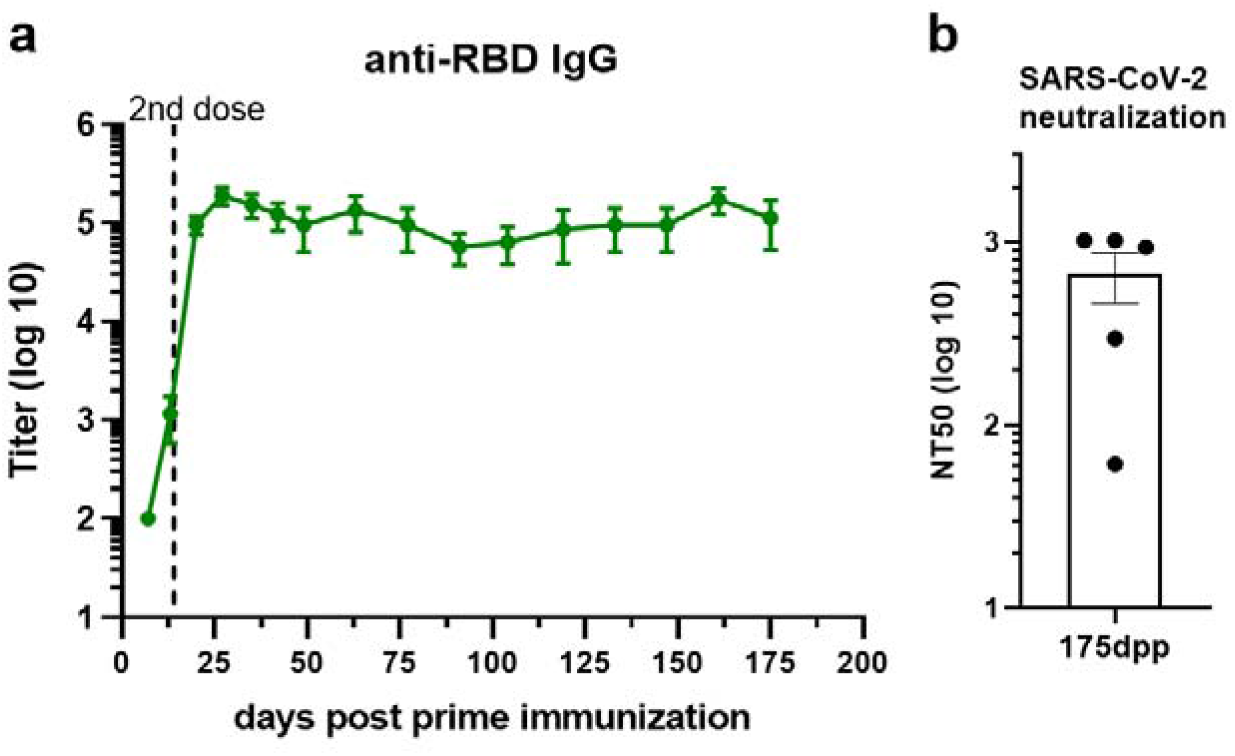
RBD+Alum+U-Omp19 immunization induces a long-term antibody response. **A**. Kinetics of RBD-specific IgG endpoint titers in sera of RBD+Alum+U-Omp19 BALB/c immunized animals by ELISA. Points are means ± SEM. **B**. Bar plot of neutralizing-antibody titer against wild-type SARS-CoV-2 at day 175 post prime immunization (dpp). Neutralization titer was defined as the serum dilution that reduces 50% the cytopathic effect (NT50). Data are mean ± SEM.

### U-Omp19+Ag+Alum formulation induces neutralizing Abs against multiple SARS-CoV-2 variants

To adequately address the public health impact that newly emerging COVID-19 variants present, there is a need for vaccine-elicited antibodies that can cross-neutralize different SARS-CoV-2 variants. Thus, neutralization activity of sera against prevalent circulating variants of SARS-CoV-2 in our region: alpha (B.1.1.7, first identified in UK), gamma (P.1, first identified in Manaos, Brazil) and lambda (C.37, first identified in Peru) was evaluated and compared to neutralizing activity against wild-type reference strain. In particular, gamma and lambda variants have been shown to partially escape neutralization by antibodies triggered by previously circulating variants or vaccine induced antibodies^25,26^. Noteworthy, antibodies induced after vaccination with the formulation containing U-Omp19 not only neutralize the wild-type virus but also cross neutralized alpha, lambda and gamma variants (**Fig. 5**). In contrast, antibodies produced by mice immunized with RBD+Alum could neutralize the wild-type SARS-CoV-2 and alpha variant but showed significantly lower neutralizing activity against gamma and lambda variants (**Fig. 5**). AddaVax and AddaS03 adjuvanted formulations induced similar neutralizing antibody titers against the wild-type, gamma and lambda variants (**Fig. 5**).

**Figure 5.**
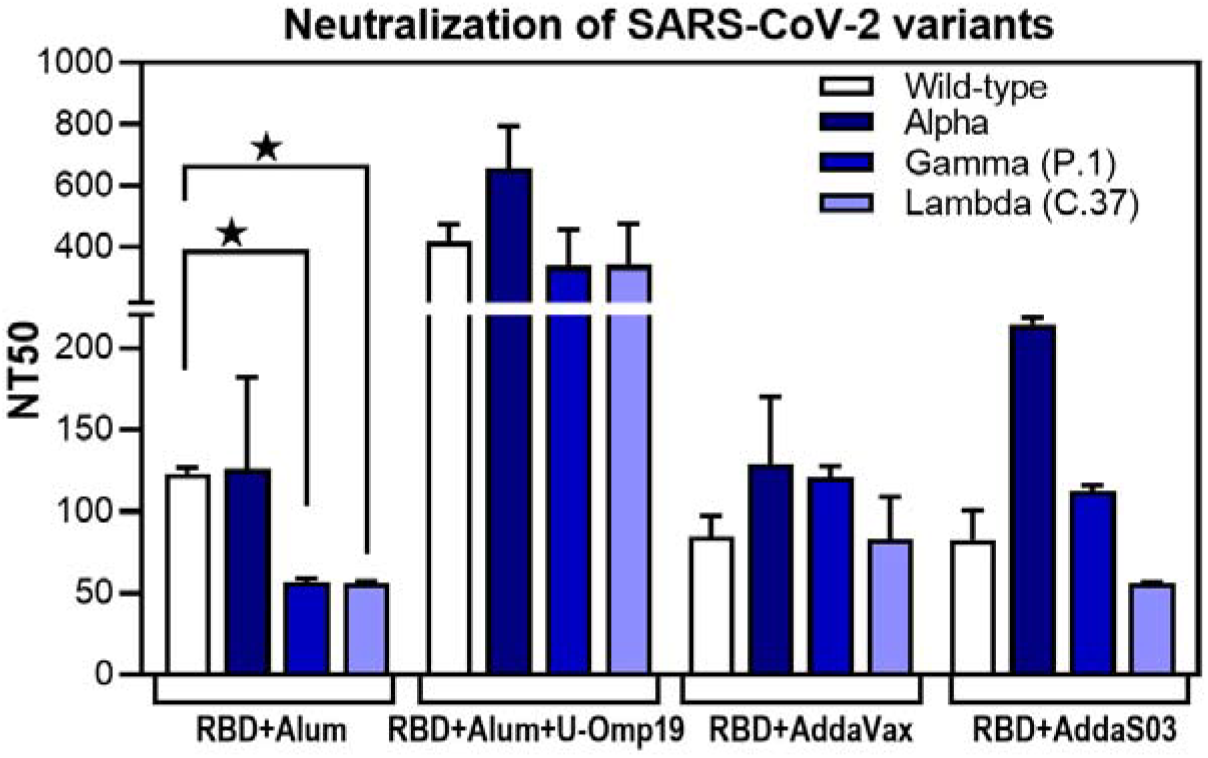
U-Omp19+Ag+Alum formulation induces neutralizing Abs against multiple SARS-CoV-2 variants. BALB/c mice were vaccinated as described in Fig. 2. Neutralizing-antibody titers against ancestral (wild-type) SARS-CoV-2 and alpha, gamma (P.1) and lambda (C.37) variants were assessed one month post second dose. Neutralization titer was defined as the serum dilution that reduces 50% the cytopathic effect (NT50). Bars represent means ± SEM. *p<0.05. T test.

Altogether these results demonstrate that addition of U-Omp19 to the Alum plus RBD vaccine formulation increases virus neutralizing antibodies, specific IgA in BAL and neutralizing antibodies of the virus in BAL. Neutralizing antibodies are proposed as the best correlate of protection thus we focused the next studies on the vaccine formulation containing U-Omp19 as adjuvant.

### U-Omp19+RBD+Alum formulation induces Ag-specific Th1 and CD8^+^ T cells in spleen and lung

In addition to memory B cells and neutralizing antibodies, induction of specific T cell immune responses could have a role in protection against SARS-CoV-2 infection^27^.

To determine T cell-mediated immune responses, splenocytes and lung cells from RBD+Alum or RBD+Alum+U-Omp19 immunized mice were stimulated with RBD or medium alone and then cytokines levels in the supernatants were measured. Both formulations were able to induce Ag-specific cytokine secretion at spleen (**Fig. 6A**). Importantly, the levels of interferon (IFN)-y were higher than interleukin (IL)-5 at spleens of both vaccine formulations. In lungs, immunization with RBD+Alum+U-Omp19 promoted a significant increment in IFN-y secretion compared with the formulation containing RBD+Alum (**Fig. 6B**). However, the Alum-adjuvanted vaccine elicited a higher amount of IL-5 by lung (**Fig. 6B**). These results suggest that U-Omp19 as adjuvant promotes a specific T cell response biased to a Th1 profile in the lung.

**Figure 6.**
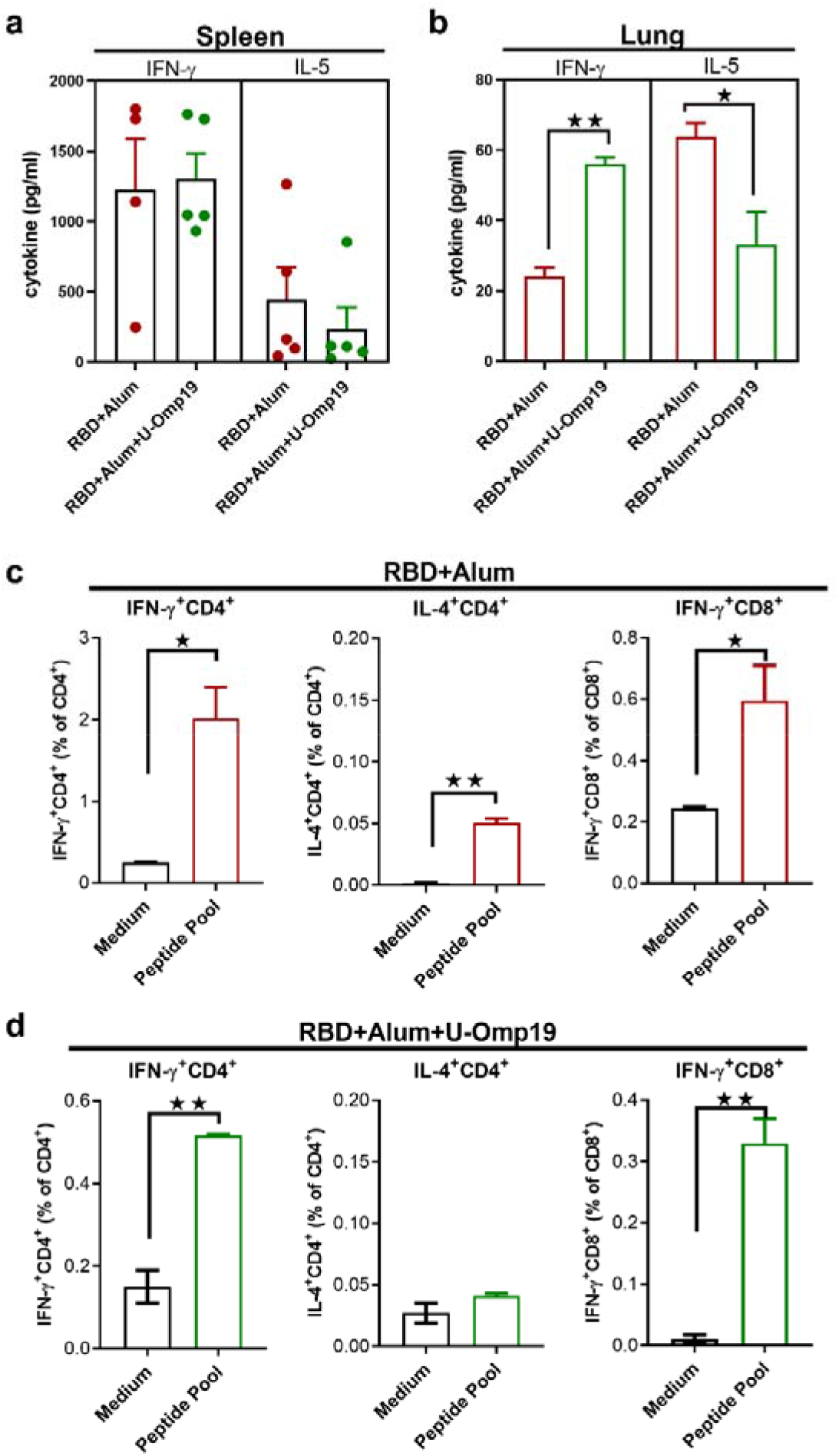
U-Omp19+RBD+Alum formulation induces systemic and mucosal Ag-specific Th1 and CD8^+^ T cells. BALB/c mice were vaccinated as described in Fig. 2. Mice were sacrificed 42 days after the first immunization to obtain spleens and lungs and T cell response was evaluated. Levels of secreted IFN-y and IL-5 following splenocytes (**A**) or lung cells (**B**) stimulation with medium or recombinant RBD were determined by ELISA. Bars are means ± SEM of pg/ml of IFN-y and IL-5 after subtracting the amount in medium stimulated cells. *p<0.05, **p<0.01. T test. **C**,**D**. Intracellular flow cytometry analysis of cytokine secreting T cells. Splenocytes from groups RBD+Alum (**C**) or RBD+Alum+U-Omp19 (**D**) were stimulated with complete medium or RBD-peptides pool for 18 h and then brefeldin A was added for 5 h. Afterward, cells were harvested and stained with specific Abs anti-CD8, and anti-CD4, fixed, permeabilized, and stained intracellularly with anti–IFN-γ and anti-IL-4. Results are presented as percentage of IFN-γ or IL-4–producing T lymphocytes. Bars are means ± SEM. *p<0.05, **p<0.01 vs. medium. T test.

To further evaluate the Th1/2 balance, IFN-γ and IL-4 producing cells were measured by intracellular cytokine staining. Spleen cells from immunized mice were stimulated with a pool of SARS-CoV-2 RBD peptides to detect antigen-specific T cell responses. Percentages of IFN-γ– producing CD4^+^ and CD8^+^ T cells were increased in both groups of mice while IL-4–producing CD4^+^ T cells were only increased after RBD+Alum administration (**Fig. 6C and D**). These results support the induction of Th1 and CD8^+^ T cell immune responses after immunization with RBD adjuvanted with U-Omp19 in combination with Alum.

### U-Omp19 as adjuvant increases neutralizing antibodies in C57BL/6 mice

Immunogenicity of vaccine formulations using Alum alone or combining both adjuvants (alum and U-Omp19) with RBD as Ag was also evaluated in the C57BL/6 mouse strain. Vaccine formulations were administered following the same schedule used for BALB/c mice, two doses every 14 days.

Both vaccine formulations induced high anti-RBD IgG titers in sera (**Fig. 7A**). Remarkably, anti-RBD IgA levels in the BAL of RBD+Alum+U-Omp19 immunized mice were higher than in the RBD+Alum immunized mice (**Fig. 7B**, \**p*<0.05). There were no differences in specific IgG levels in the BAL between both groups (**Fig. 7B**).

**Figure 7.**
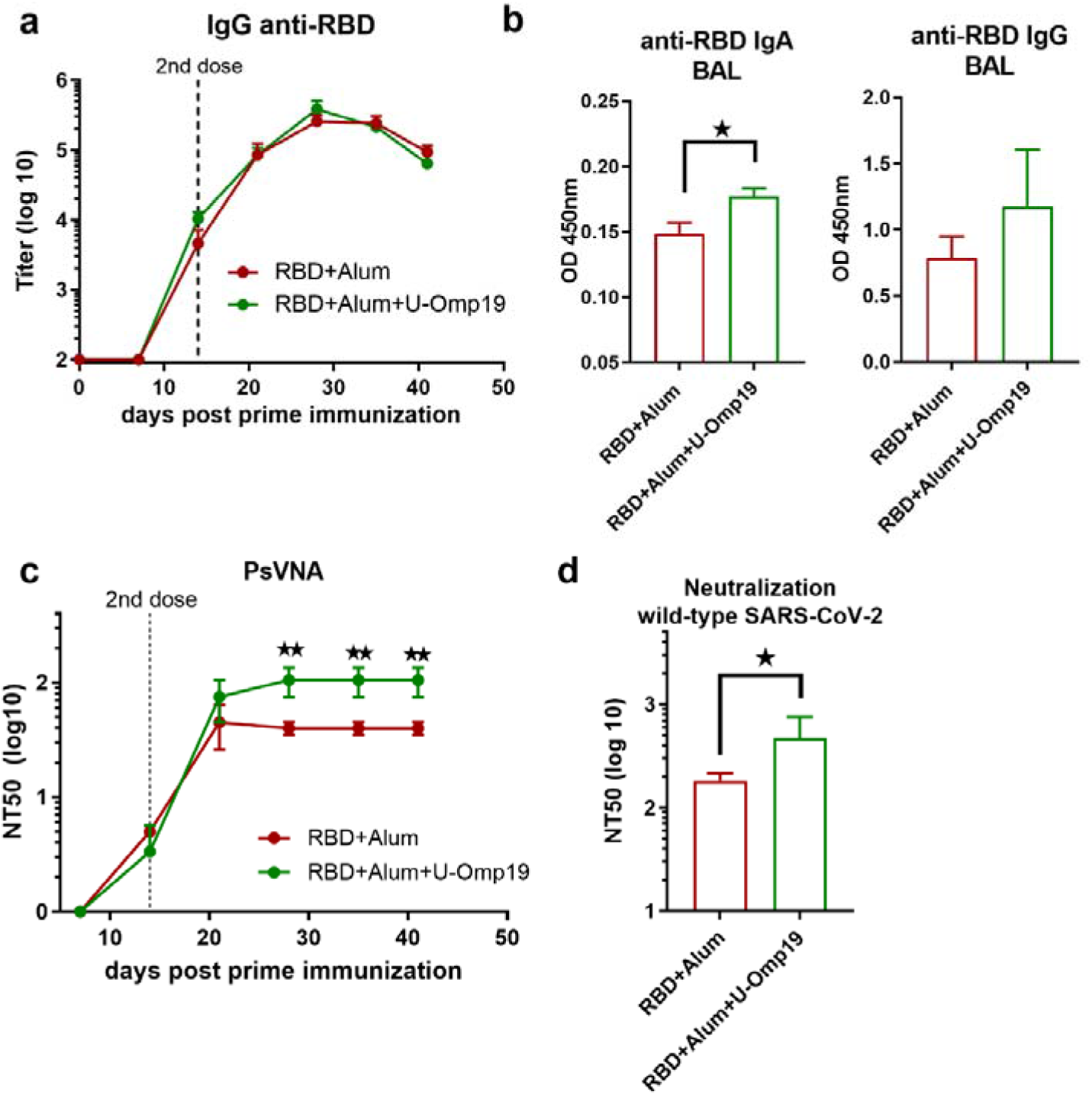
U-Omp19 as adjuvant increases neutralizing antibodies in C57BL/6. C57BL/6 mice were vaccinated at day 0 and day 14 via i.m. with RBD+Alum or RBD+Alum+U-Omp19. **A**. Kinetics of RBD-specific IgG endpoint titer in sera of immunized animals by ELISA. Points are means ± SEM. **B**. Detection of RBD-specific IgA and IgG at the bronchoalveolar lavage of immunized mice at day 42 post prime immunization. Data are optical density (OD) at 450nm. *p<0.05. Mann Whitney test. **C**. Kinetics of neutralizing-antibody titers determined by pseudo-typed SARS-CoV-2 assay. Neutralization titer was defined as the reciprocal serum dilution that causes a 50% reduction of transduction efficiency (NT50). **p<0.01. T test. **D**. Neutralization titers against wild-type SARS-CoV-2 virus at day 42. Neutralization titer was defined as the serum dilution that reduces 50% the cytopathic effect (NT50). Data are shown as means ± SEM. *p<0.05. T test.

Formulation containing RBD+U-Omp19+Alum induced higher neutralizing antibody titers than RBD+Alum formulation (**Fig. 7C**, GMT at 42 days post prime dose: 93.60 95%CI 59.99-452.5 and 37.47 95%CI 23.40-19.36 respectively).

Addition of U-Omp19 to RBD plus Alum vaccine increased the neutralizing antibody titers against authentic wild-type virus. Four weeks after second dose, sera of mice immunized with RBD+Alum+U-Omp19 produced a ten-fold increase in the viral neutralizing antibodies titer (GMT 929 95%CI 37.50-10079 **Fig. 7D**) compared with mice receiving RBD + Alum (**Fig. 7D**, GMT 99 95%CI 13.61-210.1). These results validated the data obtained in BALB/c mice and further indicate that U-Omp19 can be used under different genetic backgrounds.

### U-Omp19 adjuvanted vaccine increases RBD specific germinal centers cells and plasmablasts in the spleen

A persistent germinal center (GC) B cell response enables the generation of robust humoral immunity^28^. Therefore, specific GC B cells were evaluated in spleens from vaccinated mice one month after second dose, **Fig. 8A** shows the gating strategy used. There were no differences between groups in the total CG cells (B220^+^ CD19^+^ IgD^-^ CD95^+^ GL7^+^ cells) among spleen samples (**Fig. 8B**). Of note, there were differences in the frequency of RBD^+^ specific GC cells as mice immunized with RBD+Alum+U-Omp19 increased the percentage of RBD^+^ specific GC cells in comparison with RBD+Alum (**Fig. 8C**). Besides, the percentages of RBD^+^ specific plasma blasts (B220^+^ CD19^+^ IgD^-^ CD138^+^ cells) were also higher in animals from RBD+Alum+U-Omp19 than from RBD+Alum (**Fig. 8D**). These results indicate a better performance of the vaccine formulation containing U-Omp19 to induce specific GC and secretory B cells one month after immunization.

**Figure 8.**
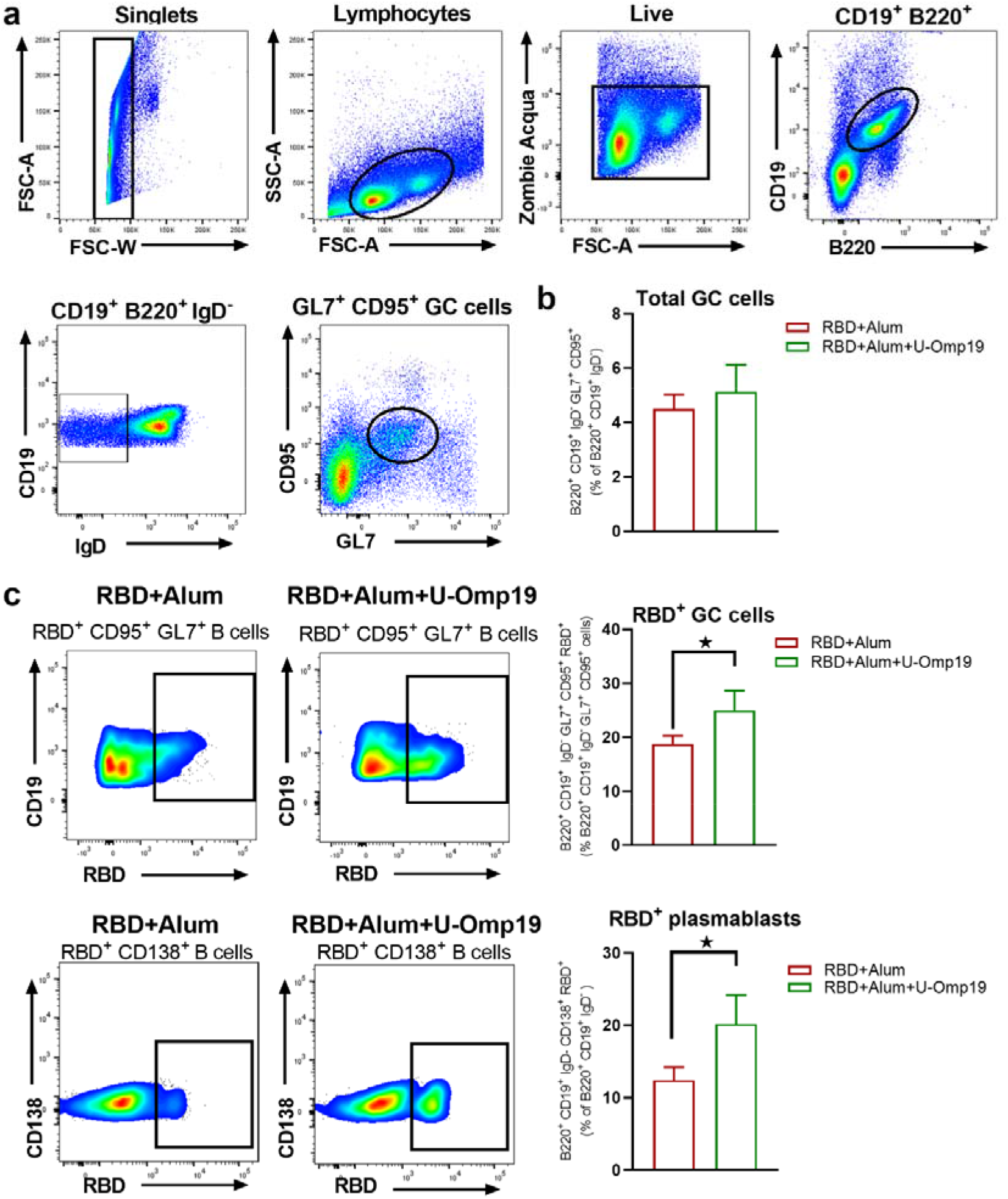
U-Omp19 as adjuvant increases RBD specific germinal centers cells and plasmablasts. Mice were vaccinated as described in Fig. 7. Flow cytometry analysis of different B cell populations at spleen of vaccinated mice were performed using, anti-CD19, anti-B220, anti-IgD, anti-CD138, anti-GL7 and anti-CD95 antibodies. Specific cells were determined by binding to fluorescent RBD. Gating strategy is shown in A. **B**. Results are presented as percentage of total GC cells (B220^+^ CD19^+^ IgD^-^ GL7^+^ CD95^+^). Bars are means ± SEM **C**. Dot plots for each group are shown (right) and results are presented as percentage of RBD-specific GC cells (B220^+^ CD19^+^ IgD^-^ GL7^+^ CD95^+^ RBD^+^). Bars are means ± SEM. \**p*<0.05. T test. **D**. Dot plots for each group are shown (right) and results are presented as percentage of RBD-specific plasmablasts (B220^+^ CD19^+^ IgD^-^ CD138^+^ RBD^+^). Bars are means ± SEM. \**p*<0.05. T test.

### RBD+Alum+U-Omp19 induces protection against intranasal SARS-CoV-2 challenge

To determine vaccine efficacy, we used a severe disease model using K-18-hACE2 transgenic mice. Infection of transgenic mice with SARS-CoV-2 results in lung disease with signs of diffuse alveolar damage, and variable spread to the central nervous system^29^. The lethal dose 50% (LD50), is estimated to be 10^4^ plaque-forming units (PFU)^30^. Vaccine formulation efficacy was evaluated in K18-hACE2 mice vaccinated with RBD+Alum+U-Omp19 or PBS (control) and challenged intranasally with 2×10^5^ PFU of SARS-CoV-2. At day 5 post infection some animals were euthanized to assess the viral load in lungs and brains. The presence of the SARS-CoV-2 virus was not detected in the lungs while very low virus titers were detected in the brains of animals vaccinated with RBD+Alum+U-Omp19 (**Fig. 9A**). In contrast a high viral load was detected in the lungs and brains of animals immunized with PBS (**Fig. 9A**). It is noteworthy that the majority of the mice vaccinated with RBD+Alum+U-Omp19 did not lose weight after challenge (**Fig. 9B**). Thus, the RBD+Alum+U-Omp19 vaccine induced protection against the experimental challenge with SARS-CoV-2.

**Figure 9.**
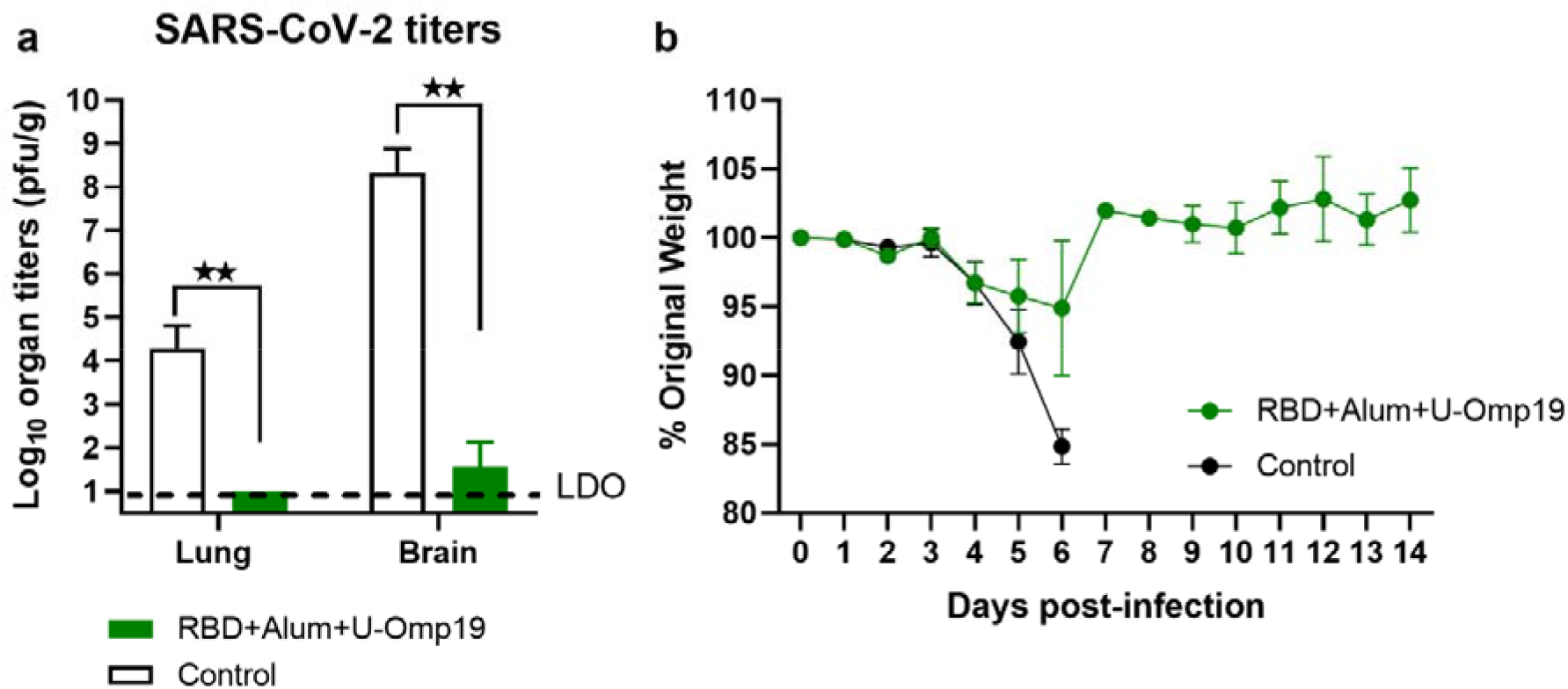
Vaccination with RBD+Alum+U-Omp19 protects K18-hACE2 transgenic mice against SARS-CoV-2 infection. Mice received PBS (Control) (n=7) or RBD+Alum+U-Omp19 (n=8) administered via i.m. route at day 0 and 14. Four weeks following immunization, K18-hACE2 mice were intranasally infected with 2 × 10^5^ PFU of SARS-CoV-2. Five days after infection lungs and brains (n=3) were obtained from groups of mice and SARS-CoV-2 virus was titrated. Bars represent the mean ± SEM. Dotted line: limit of detection (LOD). **p<0.01. T test. (**B**) Weight loss outcomes in K18-hACE2 transgenic mice vaccinated and challenge with SARS-CoV-2. Weight changes in mice were monitored daily until day 14 after infection. Points are means ± SEM of percentage of original weight.

## Discussion

There is an urgent need to develop safe, effective, and affordable COVID-19 vaccines for low-and middle-income countries. Such vaccines should rely on proven technologies such as recombinant protein–based vaccines to facilitate their transfer to emerging market vaccine manufacturers. Protein-based vaccines are classic vaccine platforms and are considered a very safe vaccine strategy. The majority of anti-viral vaccines being licensed for human use are protein-based vaccines, such as hepatitis B and human papillomavirus subunit vaccines, which have been widely administered, and present an exceptional safety profile^31^.

Our group has been working on the development of a recombinant protein–based vaccine to prevent COVID-19. We selected the SARS-CoV-2 spike protein RBD as an immunogen since it offers advantages for rational vaccine design both immunologically and from a manufacturability point of view^32^. RBD targeted binding antibodies correlate very strongly with virus-neutralizing activity in natural infections and vaccinations^33^. Therefore, selection of RBD as antigen may induce a higher proportion of neutralizing antibodies, compared to full length Spike protein immunization. Indeed, B cell repertoire analysis after RBD immunization has been shown to induce a higher proportion of neutralizing antibodies than full length Spike protein immunization^34^. Moreover, RBD binding antibodies account for more than 90% of the neutralizing activity in COVID-19 convalescent sera and vaccinated individuals^33,35^. Previous studies have found that both SARS-CoV and MERS-CoV display antibody-dependent enhancement (ADE)^36^, where non-neutralizing antibodies produced in response to a vaccine mediate virus infection via the fragment crystallizable (Fc) receptor and thus increase the risk of vaccinations enhancing viral infection^37^. Although ADE has not been reported for the existing COVID-19 vaccines, a recent study has shown that antibodies against the S protein N-terminal domain enhanced the binding capacity of S protein to ACE2 and infectivity of SARS-CoV-2^38^. To mitigate the ADE effect, minimizing non-neutralizing epitopes we decided to work with the SARS-CoV-2 spike protein RBD. Furthermore, in our hands (data not shown) and as described by others^39^, yields of recombinant RBD were much higher than those of full-length Spike, an important factor in delivering the vaccine to global population.

Different cell types have been used to produce RBD antigens, such as yeast, plant and insect cells^32,40-43^. However, production in mammalian cells^11,44-48^ may produce a RBD antigenic domain that more closely resembles that generated during virus infection in human cells (including post-translational modifications such as glycosylation and correct folding)^49^. Our results showing binding to hACE2 expressing cells and the induction of neutralizing antibody titers confirm the preservation of the RBD structure and the suitable exposition of the receptor binding motif (RBM).

In this work we have assessed the immunogenicity in mice of four different formulations containing the SARS-CoV-2 spike protein RBD with: i) human approved vaccine adjuvants: Alhydrogel (Alum), an α-tocopherol and squalene-based containing oil-in-water emulsion (AddaS03) or a squalene-based oil-in-water nano-emulsion (AddaVax) or ii) a combination of Alum and a novel adjuvant called U-Omp19, a bacterial protease inhibitor with adjuvant properties. All adjuvanted formulations induced robust anti-SARS-CoV-2 antibody responses. However, only alum containing formulations resulted in detectable antibody titers after the first immunization. This result is in line with literature reporting low immunogenicity in mice after a single dose when RBD is formulated with AddaVax^50-53^, but significant seroconversion titers after single dose when formulated with alum^40,53^ or after two doses when formulated with AddaVax or AddaS03^39,46,50,51,53^. The overall response was dominated by the IgG1 subclass in all immunized groups. Interestingly, in COVID-19 recovered individuals spike specific IgG1 antibodies correlated most closely with *in vitro* viral neutralization than other IgG subclasses^54^.

RBD-immunization with Alum, AddaS03, AddaVax or U-Omp19 + Alum induced substantial neutralizing antibody titers against Spike-pseudotyped virions and wild-type SARS-CoV-2. Notably, the formulation adjuvanted with Alum + U-Omp19 induced stronger neutralizing antibody titers when compared to formulations adjuvanted with Alum, AddaVax or AddaS03. A similar effect of U-Omp19 in antibody functionality was seen with recombinant subunit vaccine against *Trypanosoma cruzi*, where a formulation adjuvanted with U-Omp19 despite inducing lower antibody titers, showed strong antibody mediated lytic activity, which together with the induction of a Th1-biased immune response may account for the better elicited protection of this formulation^21^. This improvement in vantibody function by U-Omp19 addition to formulations could also be due to its protease inhibitor activity. U-Omp19 can inhibit neutrophil elastase^18^ and Kim et al. have recently shown that coadministration of a neutrophil elastase inhibitor enhances the affinity and function of antibodies induced by alum as adjuvant^55^. In the same work, Kim et al. showed that neutrophil elastase inhibitor supplementation can improve the efficacy of alum-adsorbed anti–SARS-CoV-2 vaccines by promoting the induction of IgA in the serum and mucosal secretions^55^. Of note, U-Omp19 adjuvanted group presented the highest levels of RBD specific IgA in serum and BAL and the highest neutralizing activity against SARS-CoV-2 Spike-pseudotyped virions in BAL. This local immune response in the lungs may be of vital importance in neutralizing the virus before infection establishment, since it has been suggested that IgA-mediated mucosal immunity may be a critical defense mechanism against SARS-CoV-2 that may reduce infectivity of human secretions and consequently viral transmission as well^24^. Moreover, secretory dimeric IgA found in mucosa has been shown to be a more potent SARS-CoV-2 neutralizer than serum IgA^56^.

Recently, new variants of concern (VOC) or interest (VOI) of SARS-CoV-2 have been identified worldwide, and many of them have been shown to partially escape neutralization by antibodies induced by infection with previous circulating variants or vaccines^25,26^. These variants harbor mutations in RBD and N-terminal domain (NTD) of the spike protein that could impair the neutralizing activity of vaccine-induced antibodies^4^. Therefore, we also evaluated the neutralizing activity induced by the vaccine formulations against highly circulating SARS-CoV-2 variants in our region: alpha (B.1.1.7), gamma (P1), and Lambda (C.37). Neutralizing activity of antibodies elicited by the vaccine formulation with alum alone was 2-fold lower for Gamma and Lambda SARS-CoV-2 variants compared to ancestral (D614G) and alpha SARS-CoV-2 variants, similar or higher variant escape to vaccine induced neutralizing antibodies was described for several vaccines^39,57,58^. Interestingly the AddaVax and AddaS03 adjuvanted formulations induced similar neutralizing antibody titers against the wild-type, gamma and lambda variants. These results are in line with published data for CHO cell expressed RBD formulated with AddaS03^39^. More importantly, RBD adjuvanted with U-Omp19 + Alum induced significantly higher and broader neutralizing activity.

These data agree with other works showing that adjuvants not only enhance immunogenicity, but also may have different potential to elicit neutralizing antibodies that provide a greater breadth of neutralization^59^.

Most effective vaccines generate prolonged immunity by eliciting long-lived plasma cells (LLPCs) and memory B cells (MBCs)^60^. LLPCs and MBCs with high affinity for the antigen are formed during germinal center reactions. In this study we have shown that RBD immunization with Alum+U-Omp19 induced higher levels of RBD specific GC B cells than RBD formulated with alum alone. It has been reported that RBD specific GC responses strongly correlate with neutralizing antibody production^61^. It has also been suggested that prolonged antigen availability along with continuous presentation of antigens via major histocompatibility complex class II can improve GC reactions^61^. We have previously demonstrated that U-Omp19 increases antigen half-life in antigen presenting cells^19,62^, and thus may increase GC reactions and CD4^+^ T cell activation. Also, it has been shown that neutrophil elastase inhibitor supplementation to alum formulations increases the frequency of GC B cells and the size of GC^55^, suggesting that enrichment of RBD-specific GC B cells induced by U-Omp19 could be related to its ability to inhibit neutrophil elastase. U-Omp19 ability to increase GC B cell reactions was also demonstrated in the context of a rabies vaccine, in which the addition of U-Omp19 resulted in enhanced immunogenicity through increasing dendritic cells activation and germinal center formation^63^.

Although antibodies have been shown to play a critical role in protection against coronavirus infections, the T cell response is still indispensable for virus clearance, decreasing severe illness, and prognostic recovery^64^. A study by McMahan et. al. in rhesus macaques suggested that vaccine induced memory T-cell responses contribute to protection against SARS-CoV-2, especially when antibodies work sub-optimally^14^. In this study, we have investigated the intensity and diversity of T cells elicited in the lungs and the spleens in response to vaccination with RBD formulated with Alum or with Alum + U-Omp19. Interestingly, immunization with RBD adjuvanted with Alum alone induced a Th2 biased response in the lungs and a Th1/Th2 profile in the spleens, the addition of U-Omp19 biased the response to a Th1 profile, with predominance of IFN-γ production in the lung. Both vaccine formulations induced both IFN-γ^+^ CD4^+^ as well as IFN-γ^+^ CD8^+^ T cell responses, while IL-4^+^ CD4^+^ were induced only after immunization with RBD + Alum. Similar results have been reported for a formulation containing RBD and Alum that induced a mixed Th1/Th2 immune response^40^. Interestingly, an RBD dimer vaccine formulated with AddaVax was unable to elicit a T cell response in the mouse model^65^. Our results highlight that Th1 or Th2 responses are mainly dependent on the type of adjuvant. Notably, SARS-CoV-2 enhanced immunopathology was associated with Th2-biased responses^66^. Therefore, the addition of U-Omp19 may be a way to prevent immunopathology during SARS-CoV-2 infection in vaccinated individuals.

Importantly, the *in vivo* functionality of RBD+Alum+U-Omp19 vaccine elicited immune responses was evaluated in a severe disease COVID19 murine model showing that this vaccine was able to confer protection in lungs and brains from i.n. SARS-CoV-2 challenged K18-hACE2 mice.

Vaccine formulation presented in this study can be further updated against new SARS-CoV-2 variants and be used as primary immunization and also as heterologous booster for other vaccines. Interestingly, priming with full-length Spike and then boosting with SARS-CoV-2 RBD ‘immuno-focuses’ neutralizing antibody responses to the RBD protein in mice and macaques^50^ and might represent an approach to redirect immunity against SARS-CoV-2 variants.

While RBD+Alum induces significant immune responses, it has been suggested that its immunogenicity should be increased. Different approaches have been proposed to increase its immunogenicity, such as i) expression as dimer^42,48,65,67^ or trimers^44^, ii) fusion to carrier proteins like human IgG Fc moiety^43,67^, tetanus toxoid^11^, interferon-α^67^, iii) addition of pan HLA-DR-binding epitope to enhance helper T cell responses ^67^, iv) using nanoparticles as delivery system^41,45,47,52,59^ or v) addition of immunopotentiators as CpG^41,42,46,59^, MPLA^43^, 3M-052 or a TLR-7/8 agonist^44,59^. Here we demonstrated that the addition of U-Omp19 to RBD + Alum formulation was able to increase the induction and breadth of SARS-CoV-2 neutralizing antibody responses, increase the frequency of RBD specific germinal center B cells and induce antigen specific Th1 and CD8^+^ T cells.

Together our results highlight that the addition of U-Omp19 could be another approach to improve vaccine formulations comprising an antigen and alum.

## Materials and Methods

### Antigen expression and purification

Codon optimized RBD containing spike signal peptide (residues 1-14) fused to the RBD domain (residues 319-541) and a C-terminal 6xHis tag was obtained. The protein was expressed from a pcDNA 3.1 plasmid in HEK 293 cells. Cells grown in monolayer were transfected with polyethylenimine (PEI) and three days after transfection the supernatant was harvested and clarified by centrifugation at 1500 x g for 15 min. The recombinant proteins were purified from supernatants by affinity chromatography with a Ni-agarose column (HisTrapTM HP, GE Healthcare, Chicago, IL), dialyzed against PBS, quantified, and stored at −80°C. LPS contamination from RBD was adsorbed with Sepharose–polymyxin B (Sigma Aldrich. St Louis, MO). Endotoxin determination was performed with a Limulus amebocyte chromogenic assay (Lonza, Basel.).

### SDS-PAGE and western blot analysis

RBD samples were run under reducing conditions by SDS-PAGE. Samples were mixed with Laemmli sample buffer with β-Mercaptoethanol. The samples were incubated at 95 °C for 5 min. Protein bands were visualized by staining with Coomassie blue R250. Bands were then transferred to a nitrocellulose membrane (GE, Healthcare. Chicago, IL), blocked with tris buffered saline (TBS)-Tween 0.05%, and incubated with human convalescent serum (1/100 dilution). An anti-human IRDye 800 (1/2000 dilution) was used as a secondary antibody for Infrared fluorescence detection on the Odyssey Imaging System.

### Size exclusion chromatography analysis of the Spike RBD domain

SEC runs were performed on a Superdex 75 column by injecting 150 µg of RBD protein in 20mM Sodium phosphate buffer, pH 7.0 and 0.2M NaCl in the absence of DTT. Runs were performed at a flow rate of 0.4 ml/min. Apparent molecular weight was calculated by calibrating the column with molecular weight markers: Bovine Serum Albumin (66.4 KDa), Ovalbumin (44.3 KDa), Papain (23 KDa) Ribonuclease A (13.7 KDa) and Aprotinin (6.5 KDa) with Vo = 8.1ml and Vo+Vi = 19.5ml.

### Adjuvants and vaccine formulations

Recombinant U-Omp19 was expressed in *E. coli* cells and purified as previously described in^17^. LPS contamination from RBD was adsorbed with sepharose–polymyxin B (Sigma Aldrich. St Louis, MO). Endotoxin determination was performed with a Limulus amebocyte chromogenic assay (Lonza, Basel). U-Omp19 preparations used contained <0.1 endotoxin units per milligram protein.

AddaVax or AddaS03 were purchased at Invivogen and Alhydrogel 2% was kindly provided by CRODA. In vaccine formulations Ag and U-Omp19 were absorbed to Alhydrogel. RBD and U-Omp19 proteins were adsorbed to Alhydrogel® (CRODA, Inc.). Protein adsorption was analyzed by sodium dodecyl sulfate-polyacrylamide gel electrophoresis, followed by Coomassie staining. Protein concentration was determined by the bicinchoninic acid method.

### Ethics statement

All experimental protocols with animals were conducted in strict accordance with international ethical standards for animal experimentation (Helsinki Declaration and its amendments, Amsterdam Protocol of welfare and animal protection and National Institutes of Health, USA NIH, guidelines). The protocols performed were also approved by the Institutional Committee for the use and care of experimental animals (CICUAE) from National University of San Martin (UNSAM) (01/2020).

### Animals and immunizations

Eight-week-old female BALB/c or C57BL/6 mice were obtained from IIB UNSAM animal facility. Animals were intramuscularly (i.m) inoculated at day 0 and 14 with i) RBD + Alhydrogel (n=5), ii) RBD + Alhydrogel + U-Omp19 (n=5), iii) RBD + AddaVax (n=5) and iv) RBD + AddaS03 (n=4). Blood samples were collected weekly to measure total and neutralizing antibody titers. At day 42 post prime immunization animals were sacrificed and spleens, lungs and bronchoalveolar lavages (BAL) were obtained.

### Determination of antibody levels in serum and BAL

RBD-specific antibody responses (IgA, IgG, IgG1, IgG2a) were evaluated by indirect ELISA. 96-well plates were coated with 0.1 µg/well of RBD in phosphate buffered saline (PBS) overnight at 4 °C. Plates were washed with PBS-Tween 0.05% and blocked with PBS-Tween 0.01% 1% non-fat milk for 1 h. Plates were then incubated with sera or BAL (diluted in PBS Tween 0.01% containing 1% non-fat milk) for 1 h and then plates were washed and incubated with HRP conjugated anti-mouse IgA, IgG (SIGMA, St. Louis, MO, USA), IgG1 or IgG2a (ThermofisherScientific, Waltham, MA) for 1 h at 37 °C. Then, TMB (3,3, 5,5 -tetramethylbenzidine) was added and reaction was stopped with H_2_SO_4_ 4 N and immediately read at 450 nm to collect end point ELISA data. End-point cut-off values for serum titer determination were calculated as the mean specific optical density (OD) plus 3 standard deviations (SD) from sera of saline immunized mice and titers were established as the reciprocal of the last dilution yielding an OD higher than the cut-off.

### Plasmids

Plasmid pCMV14-3X-Flag-SARS-CoV-2 S was a gift from Zhaohui Qian (Addgene plasmid # 145780)^68^, psPAX2 was a gift from Didier Trono (Addgene plasmid # 12260); and pLB-GFP was a gift from Stephan Kissler (Addgene plasmid # 11619).

### Cell lines

Human embryonic kidney cell line 293T expressing the SV40 T-antigen (HEK-293T, ATCC #CRL-11268) was kindly provided by Cecilia Frecha (Instituto de Medicina Traslacional e Ingeniería Biomédica, Hospital Italiano de Buenos Aires) and maintained in complete Dulbecco’s Modified Eagle’s Medium (DMEM, Gibco) containing 10% (vol/vol) fetal bovine serum (FBS, Internegocios), 100 lU/mL penicillin, and 100 μg/mL streptomycin (Gibco). For lentivirus production, HEK-293T cells were maintained in DMEM10 containing 100 µg/ml G418 (Sigma Aldrich. St Louis, MO). African green monkey kidney cell line Vero E6 ATCC #CRL-1586 were cultured at 37°C in 5% CO2 in Dulbecco’s Modified Eagle’s high glucose medium (Sigma Aldrich) supplemented with 5% fetal bovine serum (FBS) (Sigma Aldrich).

HEK-293T cells expressing the SARS-CoV-2 receptor protein ACE2 (HEK-hACE2) were established in our laboratory by lentivirus transduction. Clonal selection of HEK-hACE2 was achieved by limit dilution and selection with hygromycin 200 µg/ml.

### SARS-CoV-2 Pseudovirus (PsV) production

For SARS-CoV-2 pseudovirus production, 5×10^6^ HEK-293T cells were seeded in complete DMEM in 10-cm dishes and incubated 24 h at 37°C and 5% CO2. Pseudoviruses were obtained by co-transfection with psPAX2, pCMV14-3xFlag SARS-CoV-2 S and pLB GFP by using polyetherimide (PEI) (1:2, DNA:PEI). The supernatants were harvested at 72 h post transfection and centrifuged at 3000 × g for 15 min at 4°C. HEPES was added then to a final concentration of 20 mM to the supernatant, which was then passed through 0.45 μm filter. When necessary, pseudovirus suspensions were concentrated by an overnight centrifugation at 3000 x g at 4 °C. PVS was titrated in HEK-hACE2 cells and stored at −70°C until use.

### Pseudovirus neutralization assay (PsVNA)

HEK-hACE2 cells were seeded in 96-well plates in DMEM10 and incubated 24h at 37°C and 5% CO2. SARS-CoV-2 PsV (500-800 focus-forming units, FFU) were preincubated with serially diluted sera for 1h at 37°C. Then, PsV-sera mixture was added to HEK-hACE2 and centrifuged at 2500 rpm for 1h at 26°C. After a 72h incubation at 37°C and 5% CO2, cultures were fixed with 4% paraformaldehyde for 20 min. Transduced cells express GFP, and neutralization titer 50 (NT50) was defined as the reciprocal serum dilution that causes a 50% reduction of transduction efficiency.

### Viruses

SARS-CoV-2 reference strain (hCoV-19/Argentina/PAIS-G0001/2020 GISAID Accession ID: EPI_ISL_499083) was obtained from Dr. Sandra Gallegos (InViV working group). SARS-CoV-2 Gamma P.1 (GISAID Accession ID: EPI_ISL_2756556) and alpha (GISAID Accession ID: EPI_ISL_2756558) were isolated in Instituto de Investigaciones Biomédicas en Retrovirus y SIDA (INBIRS, UBA-CONICET) from nasopharyngeal swabs of patients. SARS-CoV-2 Lambda C.37 (hCoV-19/Argentina/PAIS-A0612/2021 GISAID Accession ID: EPI_ISL_3320903) was isolated at INBIRS from a sample of nasopharyngeal swabs kindly transferred by Dr. Viegas and Proyecto PAIS. Virus was amplified in Vero E6 cells, and each stock was fully sequenced. Studies using SARS-CoV-2 were done in a Biosafety level 3 laboratory and the protocol was approved by the INBIRS Institutional Biosafety Committee.

### SARS-CoV-2 neutralization assay

Serum samples were heat-inactivated at 56°C for 30 min. Serial dilutions were performed and then incubated for 1 h at 37°C in the presence of SARS-CoV-2 in DMEM 2% FBS. Fifty μl of the mixtures were then added t Vero cells monolayers for an hour at 37°C (MOI=0.004). Infectious media was removed and replaced for DMEM 2% FBS. After 72 h, cells were fixed with PFA 4% (4°C 20 min) and stained with crystal violet solution in methanol. The cytopathic effect (CPE) of the virus on the cell monolayer was assessed visually, if even a minor damage to the monolayer (1-2 «plaques») was observed in the well, this well was considered as a well with a manifestation of CPE. Neutralization titer was defined as the highest serum dilution without any CPE in two of three replicable wells. Otherwise, plates were scanned for determination of media absorbance at 585 nm and non-linear curves were fitted to obtain the titer corresponding to the 50% of neutralization (NT50). Neutralization assays to compare neutralization among different SARS-CoV-2 variants (alpha, gamma and lambda) were performed in the same plate for each sample.

### Determination of T cell immune responses

Four weeks after the second dose, mice were sacrificed to study cellular responses. Intracellular cytokine determination: splenocytes were cultured (4×10^6^ cells/well) in the presence of stimulus medium (complete medium supplemented with anti-CD28 and anti-CD49d) or Ag stimuli (stimulus medium + RBD-peptides + RBD protein) for 18 h. Next, brefeldin A was added for 5 h to the samples. After that, cells were washed, fixed, permeabilized, stained, and analyzed by flow cytometry. The cells were stained with Viability dye (Zombie Acqua), anti-mouse-CD8a Alexa Fluor 488, anti-mouse-CD4 Alexa Fluor 647, anti-IL-4 Brilliant Violet 421 and anti-IFN-γ PE (Biolegend. San Diego, CA).

### Determination of Ag specific B cells

Ag-specific B cells (plasmablasts and germinal center B cells) present in the spleens were determined by flow cytometry. Splenocytes were plated (2×10^6^ cells/well) and stained with Viability dye (Zombie Acqua), anti-B220 Alexa Fluor 594, anti-CD19 APC/Cy7, anti-CD138 Brilliant Violet 785, anti-IgD Brilliant Violet 605, anti-GL7 Alexa Fluor 488 and anti-CD95 PE (Biolegend. San Diego, CA). For Ag-specific detection, cells were also stained with RBD Alexa Fluor 64. Next, cells were washed, fixed and analyzed by flow cytometry.

### Vaccine efficacy in K18-hACE2 mice

Four-week-old K18-hACE2 mice (n=7-8 per group) from Jackson Laboratory were used for evaluating vaccine efficacy. Mice were separated into two groups: i) control (n=7) PBS immunized and RBD+Alum+U-Omp19 immunized (n=8). Mice in each group included males and females. They were i.m immunized at day 0 and 14 as described for immune assays. Four weeks post second vaccination, mice were challenged intranasally (i.n.) with 10^5^ PFU of SARS-CoV-2 strain WA1/2010 in each nare. Then, they were monitored daily for weight loss and signs of disease for two weeks post-challenge. Three mice per group were euthanized at day 5 post-challenge to evaluate organ viral loads, by plaque assay on Vero E6 cells.

### Statistical analysis

Statistical analysis and plotting were performed using GraphPad Prism 8 software (GraphPad Software, San Diego, CA). In experiments with more than two groups, data were analyzed using one-way ANOVA with a Bonferroni post-test. When necessary, a logarithmic transformation was applied prior to the analysis to obtain data with a normal distribution. In experiments with two groups, an unpaired t test or Mann–Whitney U test were used. A p value <0.05 was considered significant. When bars were plotted, results were expressed as means ± SEM for each group.

## Data availability

The authors declare that the data supporting the findings of this study are available from the corresponding author upon reasonable request.

## Acknowledgements

This work was supported by grants from Agencia Nacional de Promoción de la Investigación, el Desarrollo Tecnológico y la Innovación (AGENCIA I+D+i) and Ministerio de Ciencia, Tecnología e Innovación (IP COVID-260 and FONARSEC 0001); the Bill and Melinda Gates Foundation through the Grand Challenges Explorations Initiative (OPP1119024) to J.C. and from National Institute of Allergy and Infectious Diseases of the National Institutes of Health under Award Number R01AI153433 to A.J.A.

The authors thank to the staff at the Instituto de Investigaciones Biotecnológicas and the Animal Facility of Universidad de San Martin who facilitate animal studies in this work. The authors would also like to thank Danielle Porier, Krisangel López, and Manette Tanelus for technical assistance with the SARS-CoV-2 challenge studies.

## Contributions

J.C. and K.A.P. were responsible for overall experimental design and supervision of studies. D.E.A. constructed and expressed the RBD protein, designed and supervised neutralization studies. L.M.C. designed and conducted experiments, collected data and performed data analysis. L.M.S, L.A.B and M.L.D purified RBD, formulated vaccine and conducted experiments. E.F.C. performed pseudovirus neutralization studies and data analysis. C.P.C. conducted humoral and cellular studies and data analysis. A.J.A. design, supervise and conduct animal challenge studies and data analysis. W.S. performed animal challenge studies and data analysis. J.A. conducted long-term humoral response studies. P.S.P, and I.M. performed neutralization studies with wild-type and SARS-CoV-2 variants. A.V. and M.S. isolated SARS-CoV-2 variants. L.B.C. performed RBD characterization by size exclusion chromatography. L.M.C, K.A.P. and J.C. wrote the manuscript. All authors contributed to manuscript editing.

